# Development of subcortical volumes across adolescence in males and females: A multisample study of longitudinal changes

**DOI:** 10.1101/171389

**Authors:** Megan M. Herting, Cory Johnson, Kathryn L. Mills, Nandita Vijayakumar, Meg Dennison, Chang Liu, Anne-Lise Goddings, Ronald E. Dahl, Elizabeth R. Sowell, Sarah Whittle, Nicholas B. Allen, Christian K. Tamnes

**Affiliations:** Department of Preventive Medicine, Keck School of Medicine, University of Southern California, Los Angeles, CA, USA; Department of Psychology, University of Oregon, Eugene, OR, USA; Phoenix Australia: Centre for Posttraumatic Mental Health, Department of Psychiatry, The University of Melbourne, Melbourne, AU; Institute of Child Health, University College London, London, UK; Institute of Human Development, University of California Berkeley, Berkeley, CA, USA; Department of Pediatrics, Keck School of Medicine, University of Southern California, and Children’s Hospital Los Angeles, Los Angeles, CA, USA; Melbourne Neuropsychiatry Centre, Department of Psychiatry, The University of Melbourne and Melbourne Health, Melbourne, AU; Department of Psychology, University of Oslo, Oslo, Norway

**Keywords:** neurodevelopment, subcortical, adolescence, sex differences, replication, longitudinal, Magnetic Resonance Imaging, amygdala, hippocampus

## Abstract

The developmental patterns of subcortical brain volumes in males and females observed in previous studies have been inconsistent. To help resolve these discrepancies, we examined developmental trajectories using three independent longitudinal samples of participants in the age-span of 8-22 years (total 216 participants and 467 scans). These datasets, including *Pittsburgh* (PIT; University of Pittsburgh, USA), *NeuroCognitive Development* (NCD; University of Oslo, Norway), and *Orygen Adolescent Development Study* (OADS; The University of Melbourne, Australia), span three countries and were analyzed together and in parallel using mixed-effects modeling with both generalized additive models and general linear models. For all regions and across all samples, males were found to have significantly larger volumes as compared to females, and significant sex differences were seen in age trajectories over time. However, direct comparison of sample trajectories and sex differences identified within samples were not consistent. The trajectories for the amygdala, putamen, and nucleus accumbens were most consistent between the three samples. Our results suggest that even after using similar preprocessing and analytic techniques, additional factors, such as image acquisition or sample composition may contribute to some of the discrepancies in sex specific patterns in subcortical brain changes across adolescence, and highlight region-specific variations in congruency of developmental trajectories.

## 1. Introduction

Developmental patterns of brain morphology, and sex differences in this structural variation, exist due to both global and local maturational changes (Sowell, Thompson et al. 2004, Tamnes, Walhovd et al. 2013, Erus, Battapady et al. 2015, Giedd, Raznahan et al. 2015, Narvacan, Treit et al. 2017). Determining when and how sex differences emerge in the developing brain is essential to understanding differential risk for disease, especially psychopathology (Kessler, McGonagle et al. 1993, Kessler, Berglund et al. 2005), as well as life-long sex differences in various cognitive and behavioral traits (Choudhury, Blakemore et al. 2006, Rose and Rudolph 2006, Roalf, Gur et al. 2014, Gur and Gur 2016). For example, late childhood and adolescence is a time period when many forms of psychopathology begin to emerge and do so in a sex-specific fashion, with disproportionate increases in rates of anxiety and depression seen in girls and a higher prevalence of externalizing behaviors and substance use disorders in boys (Kessler, Berglund et al. 2005, Kuhn 2015). Given that structural and functional abnormalities in subcortical regions have been associated with these various mental health problems, it is thought that plausible sex differences in the development of subcortical structures may be pertinent to explaining sex differences in onset, prevalence, and progression of mental health disorders (Paus, Keshavan et al. 2008, Gogtay and Thompson 2010, Shaw, Gogtay et al. 2010). As such, a number of sex differences have been reported in structural magnetic resonance imaging (MRI) growth trajectories of subcortical structures. However, developmental patterns observed in these structures have been inconsistent across studies, and there has yet to be a consensus as to how these patterns differ between sexes (Sowell, Trauner et al. 2002, Lenroot, Gogtay et al. 2007, Ostby, Tamnes et al. 2009, Dennison, Whittle et al. 2013, Wierenga, Langen et al. 2014, Narvacan, Treit et al. 2017).

To date, studies have reported discrepant findings including growth versus reduction of the thalamus and basal ganglia beginning in late childhood, as well as stability versus continuing growth of the amygdala and hippocampus across adolescence (Giedd, Vaituzis et al. 1996, Sowell, Trauner et al. 2002, Ostby, Tamnes et al. 2009, Koolschijn and Crone 2013, Wierenga, Langen et al. 2014). Similarly, reported sex differences in these trajectories remain variable. From a study design perspective, it is believed that longitudinal study designs that are able to better account for both within - and between - individual differences over time may help to improve our understanding of cross-sectional findings that focus on mean group differences between the sexes (Crone and Elzinga 2015). As such, longitudinal MRI studies using raw volumes (uncorrected for whole brain size or other allometric scaling) consistently show larger volumes in males as compared to females (i.e. main effects) (Dennison, Whittle et al. 2013, Raznahan, Shaw et al. 2014, Wierenga, Langen et al. 2014, Narvacan, Treit et al. 2017). However, findings are less clear in terms of sex differences in the trajectories (i.e. slopes) of development seen across childhood and adolescence. Based on using raw volume estimates (i.e. trajectories reported without including allometric scaling), some studies report sex differences in neurodevelopmental trajectories of subcortical regions (Dennison, Whittle et al. 2013, Goddings, Mills et al. 2014, Raznahan, Shaw et al. 2014), whereas other studies find no difference between the sexes (Wierenga, Langen et al. 2014, Narvacan, Treit et al. 2017).

These discrepant observations in studies of subcortical volume development and sex differences in these patterns may be due to a number of factors, including cohort effects inherent to the sample, variation in study design, image acquisition and preprocessing, and/or statistical modeling approaches. In terms of image processing, dissimilarities have been reported in the absolute volume estimates as well as in the reliability of subcortical brain structures across different freely available automated segmentation software (Morey, Selgrade et al. 2010, Makowski, Beland et al. 2017). In addition, software packages vary in their methodology for processing longitudinal scans. For example, FreeSurfer’s longitudinal pipeline includes creating an unbiased within-subject template space to help reduce random variation and improve the sensitivity of detecting changes over time (Reuter, Schmansky et al. 2012). Recently, a longitudinal cortical thickness pipeline has also been developed as part of the ANTs software (Tustison, Holbrook et al. Unpublished). To our knowledge, other commonly used software packages for structural analysis (i.e. CIVET (Zijdenbos, Forghani et al. 2002), MAGeT (Chakravarty, Steadman et al. 2013), and FSL (Zhang, Brady et al. 2001)) do not account for within-subject variance in a similar fashion during the preprocessing stream. Beyond software, differences in quality control (QC) procedures utilized across studies may also impact the results (Ducharme, Albaugh et al. 2016).

From a statistical perspective, the inclusion of covariates and/or statistical model vary widely by study and may impact results (Vijayakumar, Mills et al. Accepted). For example, during statistical testing the inclusion of a ‘global’ or ‘allometric’ covariate to account for *between subject* differences in body size or weight (Sanfilipo, Benedict et al. 2004) may directly influence sex differences that are identified (Lenroot, Gogtay et al. 2007, Dennison, Whittle et al. 2013). Moreover, despite sex differences in allometric variables (i.e. whole brain or intracranial volume), recent findings suggest that the variability of anatomical volumes are not equal between the sexes (males show larger variance expressed at both upper and lower extremities of the distributions) (Wierenga, Sexton et al. 2017), allometric covariates follow non-linear developmental patterns from childhood to adulthood (Mills, Goddings et al. 2016, Reardon, Clasen et al. 2016), and regions including the thalamus, striatum, and pallidum show hypoallometric scaling with whole brain size (i.e. volumes become proportionately smaller with increasing head size) (Reardon, Clasen et al. 2016). Moreover, the inclusion of an allometric term may be redundant when examining longitudinal change using hierarchical modeling, as each subject receives its own intercept and slope (Crone and Elzinga 2015). Thus, the *between-subject* variance due to individual differences in head size is captured at the individual level over time; allowing for better characterization of changes in regional volume estimates over time.

Study results may vary based on the type of statistical analytic techniques employed. Although longitudinal studies have typically used linear mixed effect modeling (LME) to describe age-related changes, the model terms are diverse (Vijayakumar, Mills et al. Accepted). For example, studies have differed in their modeling approach, including use of polynomial terms (e.g. quadratic or cubic), model selection strategy (e.g. top-down or likelihood indices), testing males and females separately and/or including sex as an interaction term, as well as regarding the inclusion of other confounding factors (Ruigrok, Salimi-Khorshidi et al. 2014). Moreover, while LME including polynomial terms remains a popular approach, polynomials are rather restrictive, whereas other modeling techniques, such as general additive modeling (GAMM), may allow for a more flexible fit of a curve to the data. Specifically, GAMM replaces the linear slope parameters with ‘smooth’ functions to find the optimal functional form between the predictor and response (Jones and Almond 1992). Given the existing discrepancies in the existing literature and the vast array of methodology (including software, quality checking procedures, and model terms) utilized between studies, there remains an important gap in our knowledge regarding the reproducibility of possible sex differences in subcortical neurodevelopmental trajectories across childhood and adolescence.

The goal of the current study was to utilize identical image processing and analysis methods in three independent longitudinal neuroimaging samples to describe the development of uncorrected subcortical volumes for males and females from late childhood into young adulthood. This study is part of an international collaboration project intended to improve the reliability and efficiency of neurodevelopmental research by simultaneously analyzing multiple existing neuroimaging datasets (Mills, Goddings et al. 2016, Tamnes, Herting et al. 2017). By keeping longitudinal preprocessing methods, quality control procedures, and statistical methods constant across samples, we can assess and interpret the potential impact of sample and acquisition differences on brain development patterns in males and females. Moreover, given inherent study design differences between the longitudinal samples (e.g. age ranges and scan follow-up), we explored age and age by sex relationships in each sample using both the more flexible general additive modeling (GAMM) approach as well as the more common general mixed-effects modeling (LME). Because LME is the most commonly used approach in longitudinal MRI studies (Vijayakumar, Mills et al. Accepted), LME estimates in the current study were included in order to help directly compare our results with those reported in previous studies. Thus, we aimed to examine the consistency and reproducibility of neurodevelopmental change for subcortical gray matter regions, including the thalamus, caudate, putamen, pallidum, hippocampus, amygdala, and nucleus accumbens in males and females.

## 2. Materials and Methods

### 2.1 Participants

This study analyzed data from typically developing youth from three separate cohorts collected utilizing longitudinal designs at three separate sites in independent research projects: *Pittsburgh* (PIT; University of Pittsburgh, USA), *NeuroCognitive Development* (NCD; University of Oslo, Norway), and *Orygen Adolescent Development Study* (OADS; The University of Melbourne, Australia). Each project was approved by their respective local review board and informed consent/assent was obtained from parents and children prior to data collection. In order to best account for within-subject variance, only participants with ≥2 scans from each cohort were included in analyses. Details regarding participant recruitment in each project have been previously described (Yap, Allen et al. 2011, Tamnes, Walhovd et al. 2013, Herting, Gautam et al. 2014). By study design, all projects enrolled typically developing children and adolescents at baseline, although OADS over-sampled children at both high and low temperamental risk of developing psychopathology. Only data passing QC procedures from typically developing youth were included in the current study. Demographic information and sample distributions for each sample are presented in Table 1 and Figure 1. For the PIT dataset, 126 participants were recruited and scanned at baseline, with 20 not completing their follow-up visit, and 33 excluded due to poor image quality of the MRI (see additional details of QC procedures in section 2.2). For NCD, 111 participants were recruited and scanned at baseline; 26 were unable to complete their follow-up visit, and 9 were excluded due to poor image quality. For OADS, 177 participants completed a baseline visit, of which 45 did not complete any additional follow-up visits, 61 were excluded due to psychiatric history or medical illness and 4 were excluded due to poor image quality. The final samples thus included 73 participants from PIT, 76 from NCD, and 67 from OADS. In total, the present study included 216 participants (110 females) and 467 scans covering the age range of 8 to 22 years.

**Figure 1.**
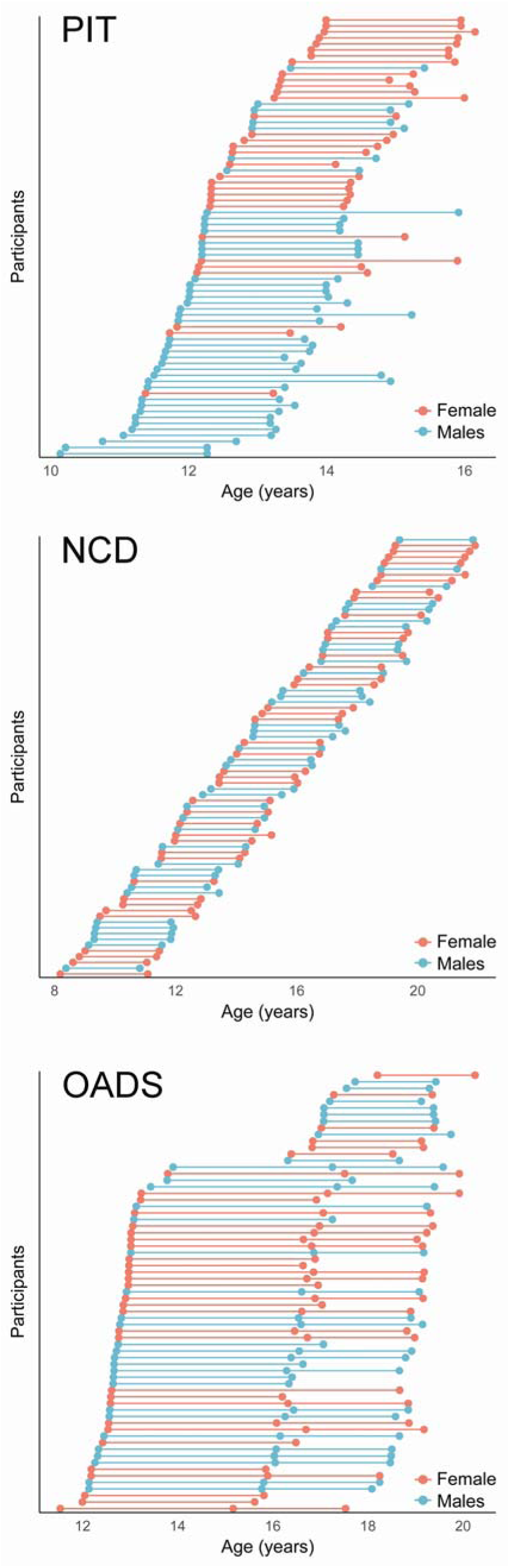
Age and sex distributions for each sample.

### 2.2 Image Acquisition and Analysis

T1-weighted anatomical scans were obtained at the three sites using different MRI scanners and sequences (see Supplementary Material). At each site, a radiologist reviewed all scans for incidental findings of gross abnormalities. Image processing, including whole brain segmentation with automated labeling of different neuroanatomical structures, was performed using the longitudinal pipeline of FreeSurfer 5.3 (http://surfer.nmr.mgh.harvard.edu; (Fischl, Salat et al. 2002, Reuter, Schmansky et al. 2012). The longitudinal pipeline includes creating an unbiased within-subject template space and image using inverse consistent registration. Skull stripping, Talariach transform and atlas registration, and parcellations are initialized in the common within-subject template, which increases reliability and statistical power. Similar standard QC procedures were carried out between sites. QC details were as follows: 1) all raw images were visually inspected for motion prior to processing, 2) post-processed images were visually inspected by trained operators for accuracy of subcortical segmentation by the longitudinal pipeline for each scan per participant, 3) images with inaccurate segmentation were excluded (number of participants excluded during QC is outlined above in section 2.1). No manual edits were made to subcortical regions of interests. Regions of interest for the present study included the thalamus, caudate, putamen, pallidum, amygdala, hippocampus, and nucleus accumbens for each hemisphere.

### 2.3 Statistical Analyses

Given previous findings highlighting hemispheric differences (Dennison, Whittle et al. 2013, Herting, Gautam et al. 2014), we first examined if patterns of change differed by hemisphere by plotting LOESS (locally weighted scatterplot smoothing) curves to each dataset. Overall, trajectories were similar between hemispheres (see Supplementary Material SFigures 1-7), and therefore left and right hemisphere volume estimates were averaged for all subsequent analyses. Given that one of the primary aims of the study was to examine the distinct developmental trajectories in males and females, and preliminary exploratory LOESS plots confirmed the shape of developmental trajectories varied between males and females (see Supplementary Material SFigures 1-7).

To more fully understand sex differences in subcortical volume changes from late childhood and throughout adolescence, analyses were performed to examine age, sex, and age by sex relationships both together across sites, as well as in each sample separately, using mixed-effects modeling with both generalized additive models (GAMM) (mgcv package version 1.8-17) and general linear mixed effects modeling (LME) (R version 3.4.0; nlme package version 3.1-131). Follow-up analyses were also conducted to examine age trajectories in each sex separately, both based on the 3 samples as well as in each sample separately.

#### 2.3.1 GAMM

Unlike parametric general linear modeling, GAMM does not require *a priori* knowledge of the functional form for the data and is an extension of LME; rather GAMM replaces one or more of the linear predictor terms with a ‘smooth’ function term. The non-linear smooth function describes the best relationship between the covariate(s) and the outcome variable of interest. For GAMM analyses on subcortical structure volume, the main predictor was age. GAMM can be represented by the following formula:

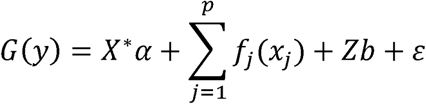

where G(y) is a monotonic differentiable link function, α is the vector of regression coefficients for the fixed parameters; X* is the fixed-effects matrix; *f_j_* is the smooth function of the covariate *x_j_*; Z is the random-effects model matrix; b is the vector of random-effects coefficients; and *ɛ* is the residual error vector.

Using this approach, GAMM models were implemented to examine age, sex, as well as an age*sex interaction to test a sex difference in the intercept (main effect of sex) as well as a sex difference in the trajectory or slope of age (age*sex interaction), respectively; while also controlling for sample at the level of intercept (main effect of sample) and slope (age*sample term). Importantly, sex was coded as a factor (male=0, female=1), allowing for each term to reflect the following: sex term reflected the difference in intercept in females as compared to males; age term reflected the slope of age for males; age*sex term reflected the difference in slope of females as compared to males. To better understand significant differences in age trajectories between the sexes, GAMM estimates for age (controlling for sample) were also implemented in each sample separately. Lastly, sample was converted to an ordered factor and GAMM models were implemented to directly test significant differences in the slopes of age between each sample for each region of interest. Thus, GAMM age, sex, and age*sex models were updated to use the previous covariates of sample and sample*age as contrasting factors. Sample was coded as a factor and two models were implemented to test sample differences: one model included sample as a factor with OADS=1, NCD=2, and PIT=3 (in order to compare OADS vs. NCD and OADS vs. PIT), and the second model included sample as a factor with NCD=1, OADS=2, and PIT=3 (in order to compare NCD vs. PIT).

#### 2.3.2 LME

LME estimates the fixed effect of measured variables on subcortical volume while including within-person variation as nested random effects in the regression model. This is done to account for individual subject effects and correlation of the data inherent to longitudinal analysis. LME can be represented by the following formula for linear changes with age both between and within participants:

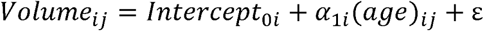

where Volume_ij_ represents the volume in an ROI at the *j*^th^ timepoint for the *i*^th^ participant, the intercept_0i_ represents the grand mean at the centered age (age 15), α_1i_ is the grand mean slope of age (linear); and ε is the residual error and reflects within-person variance. All models also included a random intercept for each participant. The linear model was then built upon to also include quadratic and cubic fixed terms to assess linear versus more complex patterns of change. The linear, quadratic, and cubic models were as follows:

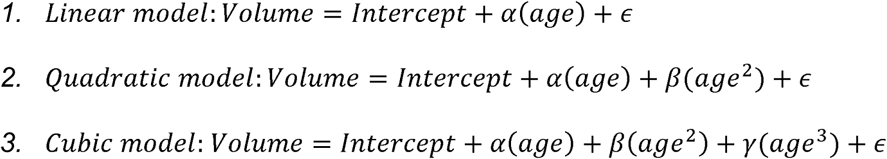

where α, β, and γ represent the effects of each fixed term. Likelihood ratio tests and Akaike Information Criterion (AIC) were used to compare the models and to determine which had the best fit. All models were tested against a null model that included only the intercept term, but not the fixed effect of age. The model with the lowest AIC that was also significantly different from the less complex model as determined by the likelihood ratio test was chosen as the best fit model (e.g. linear had to have a lower AIC and be significantly different from null; quadratic had to have a lower AIC and be significantly different from both the null and linear model).

Using LME, models of age were implemented on each sample separately to examine sex differences by including a term for the main effect of sex as well as an age*sex interaction to each model to test a sex difference in the intercept (main effect of sex) as well as a sex difference in the trajectory or slope of age (age*sex interaction), respectively. In the cases where polynomial LME best fits were different between males and females, sex difference were only tested by using the highest polynomial fit. That is, if a linear best fit was detected for females but a quadratic best fit for males, a quadratic fit was tested between sexes.

## 3. Results

### 3.1 Description of Developmental Age Trajectories using GAMM

GAMM estimates of developmental trajectories for volume for each region of interest in females and males based on the three independent samples are presented in Figure 2. GAMM models included age, sex, as well as an age*sex interaction to test a sex difference in the intercept (main effect of sex) as well as a sex difference in the trajectory or slope of age (age*sex interaction), while covarying for sample (sample and age*sample). A significant sex difference was detected for the smoothed slope of age for all seven regions of interest (Table 2). To better understand these differences, we examined age trajectories in each sex separately, while again covarying for sample. These results are presented in Table 3, and we below describe these developmental trajectories for each subcortical structure in females and males.

**Figure 2.**
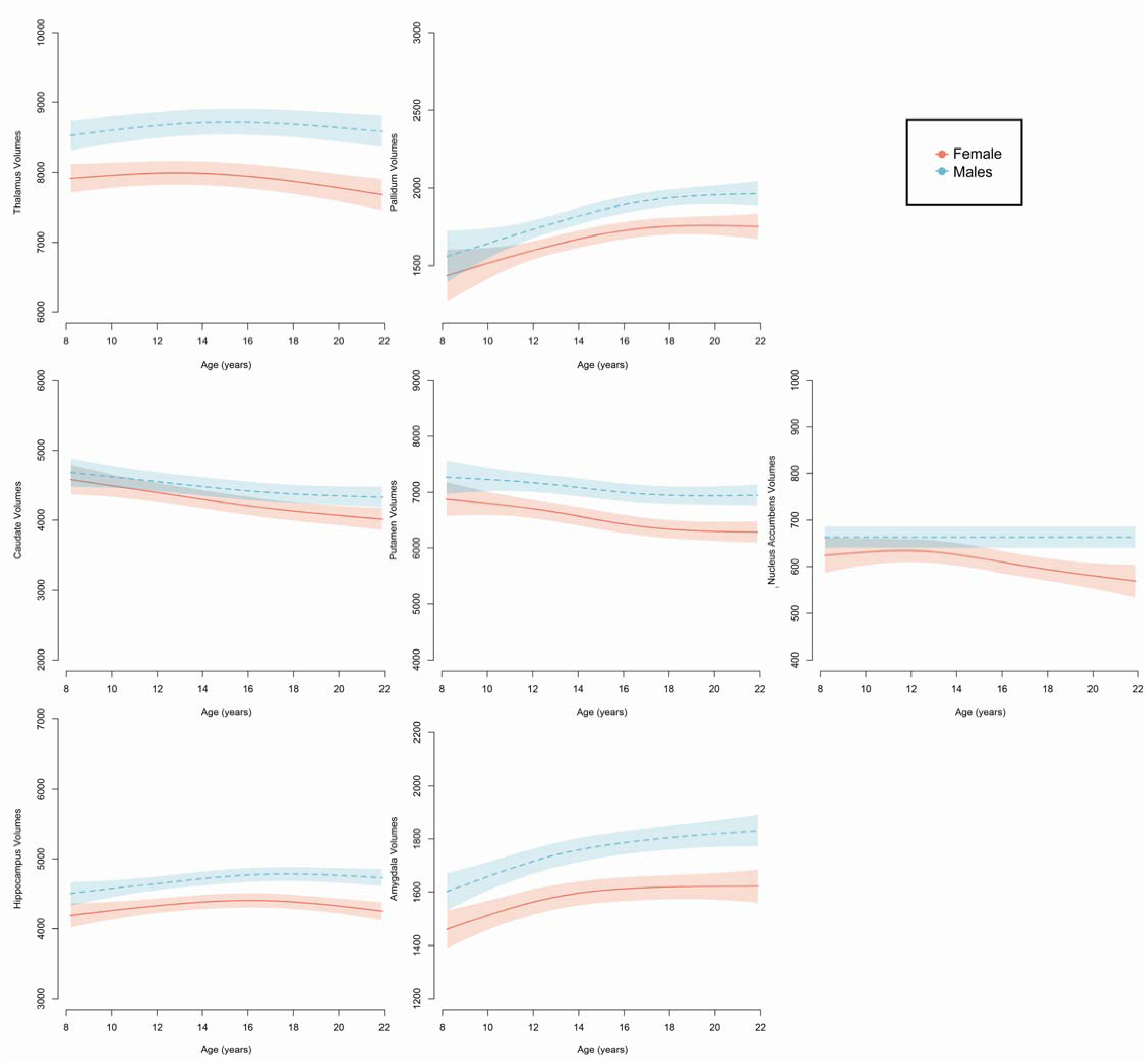
Sex differences in the developmental age trajectories for subcortical volumes based on three independent samples.

#### 3.1.1 Thalamus

Overall, females showed smaller thalamus volumes as compared to males across the entire age range of 8 to 22 years. Moreover, both males and females showed a nonlinear change with age, with decreases seen during mid-adolescence and into adulthood; however, when tested separately, the slope for age was only significant in males, and not in females.

#### 3.1.2 Pallidum

Age trajectories for each sex displayed greater divergence in pallidum volumes from ages 8 to 22 years, with males showing larger volumes as compared to females beginning in early adolescence thru young adulthood. However, when each sex was examined separately, age trajectories for the pallidum did not reach statistical significance in either females or males alone.

#### 3.1.3 Caudate

Males and females displayed similar volumes during late childhood and early adolescence, whereas females had smaller volumes compared to males by young adulthood. When each sex was examined separately, the sex difference detected in young adulthood was a result of females showing a decrease in caudate volumes across adolescence, with no significant change seen in volumes with age in males.

#### 3.1.4 Putamen

Similar to the caudate, sex differences in the putamen volumes also emerged with age, with greater sex differences seen in later adolescence and young adulthood. When examined separately, putamen volume showed a nonlinear decrease with age in females, whereas changes in volumes did not reach significance in males.

#### 3.1.5 Nucleus Accumbens

Nucleus accumbens volumes were similar in males and females from late childhood to mid-adolescence, with sex differences emerging during late adolescence and young adulthood. Examining the sexes separately revealed a significant decrease in nucleus accumbens volumes for females, with no changes in volume in males from ages 8 to 22 years.

#### 3.1.6 Hippocampus

Females showed smaller hippocampal volumes by 10 years of age compared to males. Moreover, females and males both showed significant nonlinear patterns of hippocampal growth with age, with an emergent divergence between the sexes across adolescence and into young adulthood. These nonlinear changes with age reached significant in males and females when examined separately.

#### 3.1.7 Amygdala

Females showed smaller amygdala volumes at age 8 years compared to males, with greater separation in volumes seen between the sexes with age. Again, these nonlinear changes with age reached significant in males and females when examined separately.

### 3.2. Testing Between Sample Differences

Spaghetti plots of GAMM estimates of developmental trajectories for volume for each region of interest in females and males for each of the three independent samples are presented in Figures 3-5. To directly test sample differences, previous GAMM models were updated to change the covariates of sample and sample*age as contrasting factors (sample: OADS=1, NCD=2, PIT=3). This allows for directly comparing the main effect of sample as well as if the samples have significantly different trajectories over age. GAMM estimates for each of these smooth terms are presented in Table 4 and described below.

**Figure 3.**
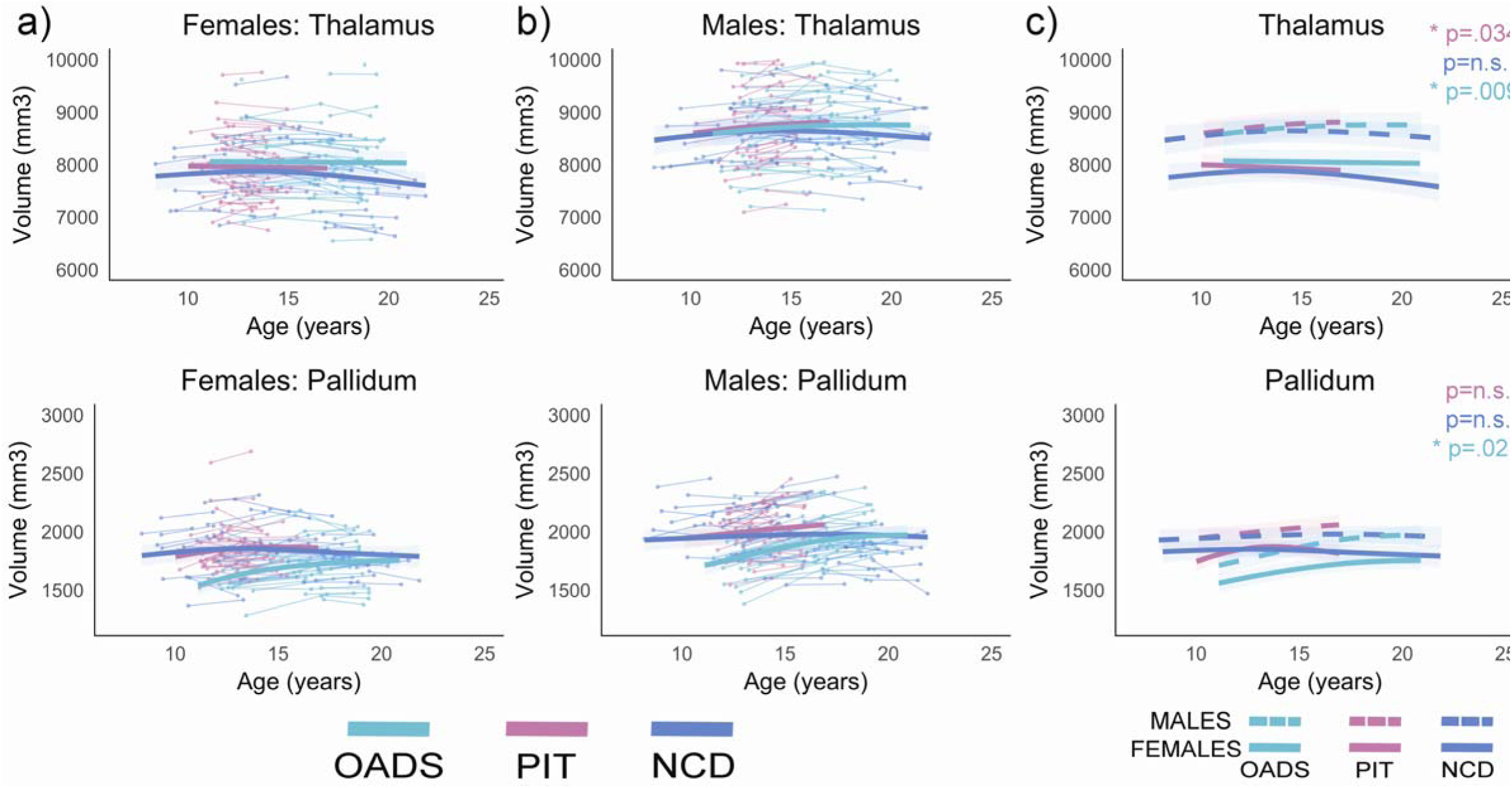
Developmental age trajectories for the thalamus and pallidum. a) Females and b) males are plotted separately. Individual datapoints are shown, connected for each participant, in the appropriate sample color. The bolded colored lines represent the GAMM fitting for each sample with 95% confidence intervals. c) Representation of GAMM fits (with 95% confidence intervals) for each sex per sample plotted together, p-values represent sex differences per sample.

**Figure 4.**
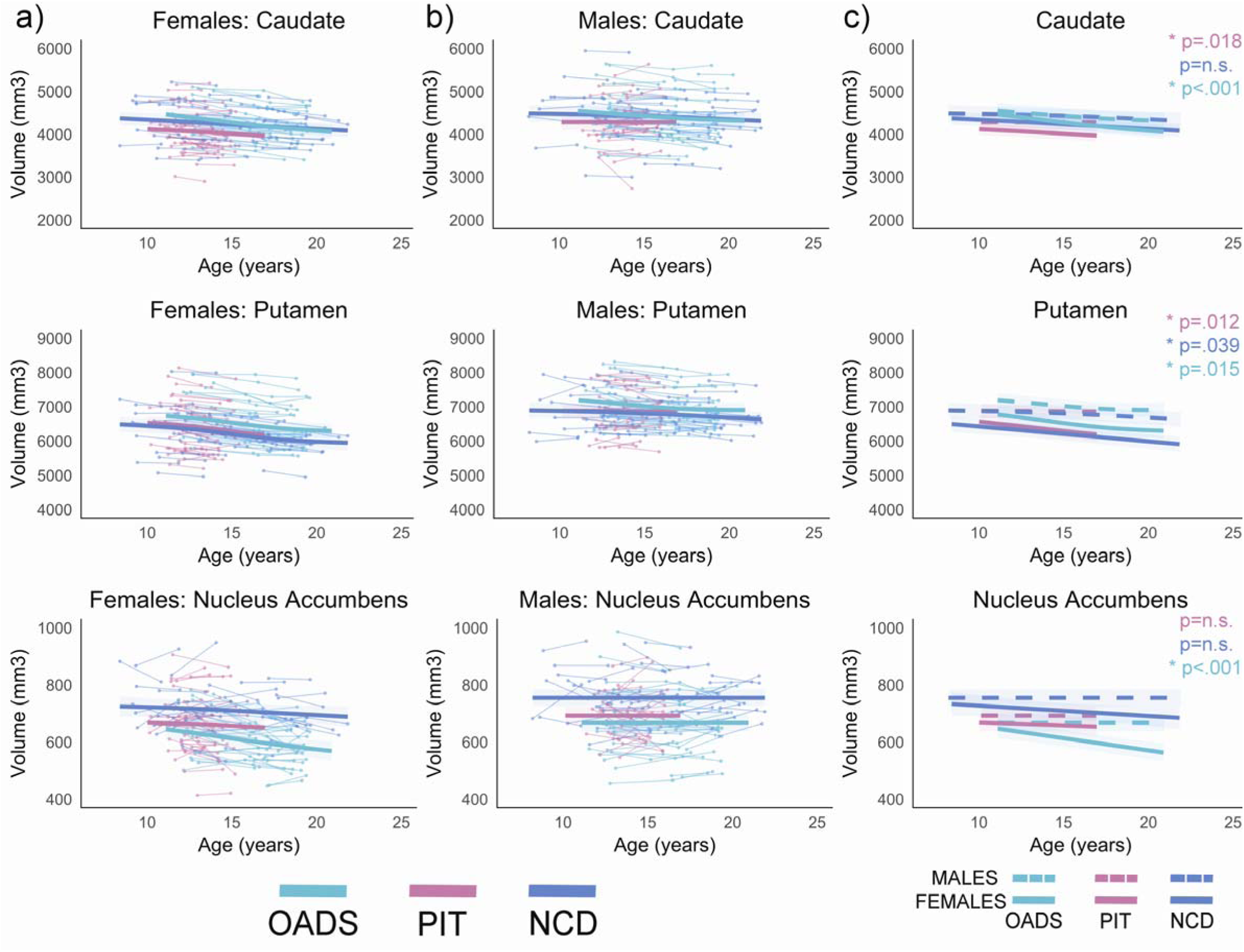
Developmental age trajectories for the caudate, putamen, and nucleus accumbens. a) Females and b) males are plotted separately. Individual datapoints are shown, connected for each participant, in the appropriate sample color. The bolded colored lines represent the GAMM fitting for each sample with 95% confidence intervals. c) Representation of GAMM fits (with 95% confidence intervals) for each sex per sample plotted together; p-values represent sex differences per sample.

**Figure 5.**
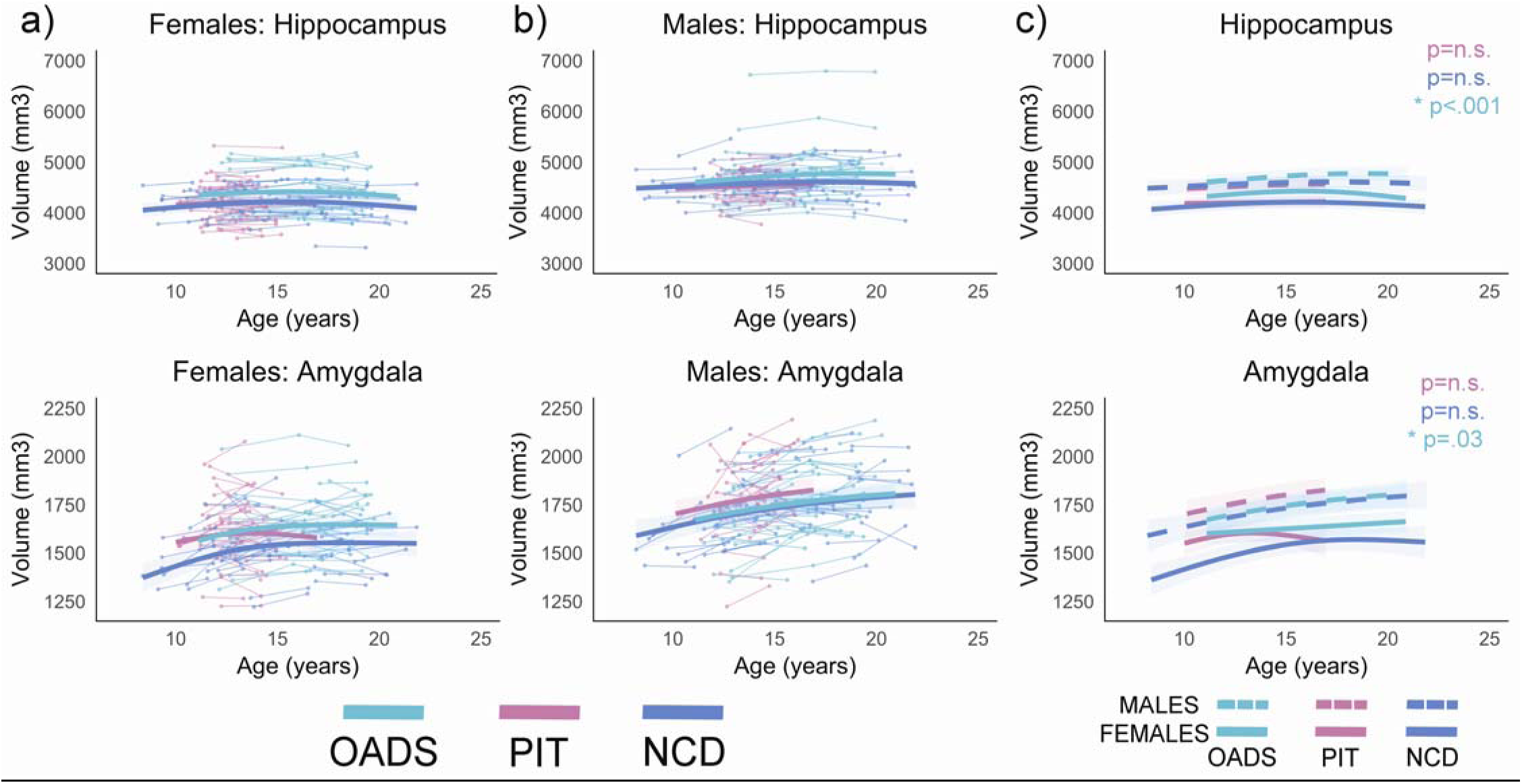
Developmental age trajectories for the hippocampus and amygdala. a) Females and b) males are plotted separately. Individual datapoints are shown, connected for each participant, in the appropriate sample color. The bolded colored lines represent the GAMM fitting for each sample with 95% confidence intervals. c) Representation of GAMM fits (with 95% confidence intervals) for each sex per sample plotted together; p-values represent sex differences per sample.

#### 3.2.1 Thalamus

The main effect of sample was not significant. However, the OADS and NCD samples showed significant sample differences in growth trajectories for the thalamus, whereas trajectories were not significantly different between PIT and OADS or NCD for this region.

#### 3.2.2 Pallidum

A main effect of sample was seen with OADS having significantly smaller volumes compared to NCD and PIT at baseline. Sample differences were also seen in age trajectories, with significant differences noted between each of the three samples (OADS vs. PIT, OADS vs. NCD, and PIT vs. NCD).

#### 3.2.3 Caudate

A main effect of sample was seen with PIT having significantly smaller volumes compared to NCD and OADS. Significant sample differences were seen in age trajectories between OADS and NCD as well as OADS and PIT samples, whereas trajectories were not significantly different between PIT and NCD.

#### 3.2.4 Putamen

A main effect of sample was seen with OADS having significantly larger volumes compared to NCD and PIT. However, the samples did not have significantly different age trajectories.

#### 3.2.5 Nucleus Accumbens

A main effect of sample was detected reflecting the largest volumes seen in NCD, followed by PIT, and OADS (all *p*‘s <0.05). However, the samples did not have significantly different age trajectories.

#### 3.2.6 Hippocampus

A main effect of sample included OADS having larger hippocampal volumes as compared to NCD and PIT. In addition, OADS and PIT samples showed significant sample differences in growth trajectories for the hippocampus, whereas trajectories were not significantly different between OADS and NCD or PIT and NCD.

#### 3.2.7 Amygdala

A main effect of sample was seen with PIT having significantly larger volumes compared to NCD, but no significant difference detected between PIT vs. OADS or NCD vs. OADS. The samples did not significantly differ for age trajectories for the amygdala.

### 3.3 Testing of Developmental Models using LME

Linear, quadratic, and cubic LME were used to determine best fit models for females and males of each sample. The highest-order polynomial model for each brain region is summarized in Table 5 (for AIC comparisons, see Supplementary Tables 1-7). LME best fits were also different between samples in both sexes for most ROIs, except for the caudate and nucleus accumbens. For the caudate, both males and females in each sample showed similar trajectories with age (PIT: linear, NCD: quadratic, and OADS: linear). For the nucleus accumbens, no significant change was found in all samples, except for OADS females, which showed a linear decrease with age. Overall, LME best fit models per sample were largely in agreement with the GAMM trajectories; the exception to this are highlighted in Supplementary Table 8 and include the pallidum for NCD females (LME=cubic, GAMM=n.s.), the hippocampus for PIT females (LME=cubic, GAMM=n.s.), the amygdala for PIT females (LME=quadratic, GAMM=n.s.) and the hippocampus for NCD males (LME=n.s., GAMM=nonlinear). Full model details for LME results when testing sex, age, and age*sex using linear and polynomial LME best fit models are presented in Supplementary Tables 10-16. Using LME, a few models that were identified as significant using GAMM did not reach significance using best fit models including the pallidum and putamen in the OADS sample and the putamen in the NCD sample (as shown in Supplementary Table 17).

## 4. Discussion

This is the first study to examine longitudinal subcortical neurodevelopmental trajectories in males and females using a multisample approach spanning ages 8 to 22 years. The current study is an extension from an on-going international collaboration project aiming to improve the understanding, reliability, and efficiency of neurodevelopmental research by simultaneously analyzing multiple existing longitudinal neuroimaging datasets (Mills, Goddings et al. 2016, Tamnes, Herting et al. 2017). By utilizing the identical longitudinal preprocessing pipeline, QC procedures, and statistical methods across samples, we aimed to shed light on the potential impact of sample and acquisition differences on our ability to detect sex differences in subcortical developmental patterns. While sex differences in patterns of subcortical development across adolescence were identified using all three datasets, divergent results were also seen for both within-sex and between-sex differences when comparing estimates from each of the independent datasets. Below we describe the findings using all samples as well as the differences detected between samples, as well as highlight the additional factors that may continue to contribute to mixed findings in our understanding of sex differences in subcortical neurodevelopment.

### 4.1 Patterns of Age Related Changes in Males and Females

GAMM estimates highlight an overall sex difference in patterns of development of subcortical volumes based on data from three independent samples (Figure 2). Significant non-linear changes were seen with age in the thalamus, curvilinear growth of the pallidum and amygdala, and decreases in the caudate, putamen, and nucleus accumbens. The average trajectories from our longitudinal datasets are largely in agreement with previous research that sensory, motor, and cognitive related subcortical regions, such as the caudate and the thalamus, undergo reduction into young adulthood (Lenroot, Gogtay et al. 2007, Raznahan, Shaw et al. 2014), as well as increases in amygdala volumes (Goddings, Mills et al. 2014).

Estimated sex differences across all participants confirmed previous findings of overall smaller volumes in females compared to males in all subcortical regions examined in the present study. In addition, significant sex differences were detected for changes with age for all regions of interest (Table 2). Of these results, replication across all three independent samples was relatively poor, as assessed by statistical results comparing age trajectory GAMM estimates. In fact, when directly testing between sample differences in age trajectories, only the putamen, nucleus accumbens, and amygdala showed no significant differences in trajectories of age development between the samples (Table 4). These findings may suggest greater generalizability of the sex differences in curvilinear amygdala growth across childhood and adolescence, with males showing significantly steeper increases compared to females. In addition, across samples, females displayed decreases in nucleus accumbens and putamen volumes, whereas males showed no change (nucleus accumbens) or less change (putamen) with age from 8 to 22 years.

### 4.2 LME versus GAMM Modeling

LME is perhaps the most commonly used statistical approaches to determining both between and within-person changes in longitudinal neuroimaging studies (Vijayakumar, Mills et al. Accepted). At the outset of the study, it was assumed that using LME might therefore allow for a more direct comparison of our results and previous studies. However, previous studies have also shown that the shape of growth trajectories can vary when examining each sex separately (Goddings, Mills et al. 2014). When using LME, this creates a challenge because in the case where the shape of the trajectory may differ between groups, putting both males and females in the same model may incorrectly assume similar shapes in growth in both sexes. For these reasons, GAMM may allow for a more flexible fit, given that is does not assume the curve to the data at the time of fitting the model. For these reasons, we chose to examine each sex separately as well as together using both LME and GAMM. Overall, strong similarities were seen in the ability for GAMM and LME modeling strategies to detect significant age-related changes in each sex separately across the three independent samples. However, when testing significant differences in changes in volumes with age between the sexes, GAMM identified changes in the pallidum for NCD and OADS and the putamen for NCD as significantly different between males and females, whereas LME models had *p*‘s>0.05. Thus, GAMM models may be able to help reframe and bring additional clarity in understanding group differences in patterns of neurodevelopment, especially when there are presumed sex differences in the shape of trajectories in males versus females.

### 4.3 Sample Consistencies and Differences

Despite our best efforts to minimize between sample effects by utilizing similar preprocessing and analytic techniques, both within-sex and between-sex trajectories of neurodevelopment were nevertheless significantly different between samples. This may suggest that factors such as population differences, sampling strategy, scanning protocols, as well as age-range and statistical power may influence best model fits in longitudinal studies of subcortical development. Because studies often differ in their age-ranges, scan intervals, and sample size, the conclusion that a particular region shows a linear, quadratic, or even cubic developmental pattern will not necessarily generalize to another study. For example, the current LME results show both linear and quadratic best fits for caudate volumes in males and females, and in contrast the putamen showed linear and cubic fits in females, but linear and quadratic for males. Similarly, previous single-sample longitudinal studies have also reported linear decreases in caudate and putamen volumes in both sexes (7-24 years; n=223 scans from 147 individuals; (Wierenga, Langen et al. 2014)); but also no change for caudate and quadratic for putamen volumes from 5 to 27 years (n=175 scans from 84 individuals; (Narvacan, Treit et al. 2017)) or quadratic for females and cubic for males from 3 to 26 years (n=829 scans from 387 individuals (Lenroot, Gogtay et al. 2007)). This poses a challenge for the field, especially given that we are rarely interested in exact ages, but rather periods of development across the life span. Furthermore, while GAMM models may provide greater precision in the description of volumetric change by moving away from more traditional polynomial assumptions in neurodevelopmental growth trajectories, significant differences were also found in the current study when using these models to examine developmental changes in subcortical volumes with age across the included samples. Given how sensitive these analyses can be to sample differences, it may be more useful as a field to focus on patterns of change (i.e. periods of relative stability/change and direction of change, as opposed to using model terms) when trying to understand overall developmental patterns, as well as providing access to statistical code in order to allow for directly testing prediction accuracy of previously published models on new datasets.

Besides inherent study population and sample differences, power is likely an issue when examining each sample separately (N’s ranging from 67-76 per study versus N=216 together). Interestingly, despite being able to better account for within-subject variability, these longitudinal findings are in agreement with recent reports that sample composition can alter age associations in large cross-sectional study designs (LeWinn, Sheridan et al. 2017). Furthermore, when examining sex differences in each sample separately (Supplementary Table 9), OADS was also found to show significant sex differences for each region of interest, whereas the other two samples were more variable. The ability for OADS to detect similar sex differences as seen with the larger combined sample may in fact be due to better within-subject estimates due to three waves of data collection as compared to the two waves design used for NCD and PIT.

Although using a commonly employed longitudinal preprocessing pipeline stream, the degree to which automated segmentation programs may contribute to the seen sample differences remains a concern. Given that manual tracing requires availability of multiple highly trained raters without intra - and inter-rater drift over time, manual tracing becomes exceeding time intensive for even medium scaled longitudinal studies that span multiple years, such as those included here (PIT=146, NCD=152, and OADS=169 scans). For these same reasons, poor segmentations are often excluded from the analyses rather than performing manual edits to the FreeSurfer subcortical volume segmentation (e.g. aseg) (as done in the current study). Thus, large scaled studies often implement automated software and in the current study we implemented the FreeSurfer longitudinal pipeline given that it was specifically created to better capture within-subject changes over time in the subcortical regions examined (as shown by intraclass correlation coefficients ranging from .90 for the left amygdala to .99 for the right caudate and putamen; note that nucleus accumbens volumes were not included in this report) (Reuter, Schmansky et al. 2012). While to our knowledge, no study has been published comparing the longitudinal pipeline estimates with manual tracing, cross-sectional studies have found that automated software tend to overestimate subcortical volumes as compare to manual tracings (Schoemaker, Buss et al. 2016, Makowski, Beland et al. 2017). Moreover, for longitudinal studies if an over estimation in volumes consistently occurs at both the between-and within-subject levels, relative differences are likely to still be meaningful (Chepkoech, Walhovd et al. 2016). However, variation in volume estimates between scanners types and acquisition parameters is likely more problematic, especially if this interacts with age. In this regard, cross-sectional studies have reported the pallidum to have poor reliability out of subcortical volume measurements using FreeSurfer, with volume estimates for this region impacted by MP-RAGE acquisition parameters, such as isotropic versus anisotropic voxel size (Wonderlick, Ziegler et al. 2009). Furthermore, low reliability of pallidum volumes, specifically, have been attributed to the T1-weighted contrast profile of the pallidum which is less distinct from its surrounding white matter as compared to other subcortical regions such as the thalamus or caudate (Fischl, Salat et al. 2002, Wonderlick, Ziegler et al. 2009). With dedication to automated software continuing to improve (e.g. FreeSurfer 6.0 was released mid-way through the current project), future studies will benefit from reductions in such potential software confounds.

### 4.4 Limitations

Recent studies document the impact of motion on structural measures (Reuter, Tisdall et al. 2015, Alexander-Bloch, Clasen et al. 2016, Ducharme, Albaugh et al. 2016), and this likely represents an especially important confound for developmental studies. We therefore conducted detailed QC of all raw and processed images and excluded participants with excessive motion. Nonetheless, future studies could benefit from employing standardized and well-documented QC procedures (Backhausen, Herting et al. 2016), and/or methods for tracking in-scanner motion, automated QC assessment, and motion correction procedures (further discussed in (Vijayakumar, Mills et al. Accepted)). Previous studies suggest that within-subject changes in puberty (both physical and hormonal) are important factors for amygdala growth in male and female adolescents (Goddings, Mills et al. 2014, Herting, Gautam et al. 2014); with very similar curvilinear amygdala growth patterns seen as reported here when raw volumes were estimated based on Tanner stage in males and females separately (Goddings, Mills et al. 2014). Puberty has also been found to relate to nucleus accumbens volumes (Goddings, Mills et al. 2014), although the trajectories do not mirror the patterns of volumetric change identified in the current study. Unfortunately, pubertal metrics were not consistent across the three cohorts; making it impossible for us to investigate how puberty may contribute to differences in amygdala and nucleus accumbens trajectories in males and females in the current study. Thus, future research is warranted to examine the contributions of hormones to sex differences in the neurodevelopmental trajectories of the amygdala and nucleus accumbens. In addition, FreeSurfer 6.0 was released mid-way through the current project, after the preprocessing for the current study was complete. Replication studies are always warranted, and should consider also examining the potential impact of the longitudinal processing stream of FreeSurfer 5.3 versus FreeSurfer 6.0, especially given the efforts of this new version on estimating the putamen.

## 5. Conclusions

Across all participants from the three independent samples, sex differences in age trajectories of volumetric development for the thalamus, pallidum, caudate, putamen, nucleus accumbens, hippocampus, and amygdala were apparent. Using a multisite approach with consistent longitudinal preprocessing (software and quality checking) and statistical analyses, generalizable patterns were found for age changes across adolescence in the amygdala, putamen, and nucleus accumbens. However, conspicuous sample differences were seen for the thalamus, pallidum, caudate, and hippocampus; perhaps a cautionary limitation when attempting to generalize subcortical findings from these regions of interests in longitudinal samples with different age ranges. Efforts aimed at improving our ability to replicate trajectories in typical development, such as the current study, are ultimately necessary in order to be able to further focus our inquiry on the factors influencing sex differences and individual differences in subcortical growth; including genetic, and/or environmental effects that may contribute to the observed differences at the group and individual-level. Furthermore, improving our ability to assess the ‘age residual’ of within-subject changes in deep gray matter structures is crucial in our ability to understand risk and resilience for psychopathology during development.

## Acknowledgments

The authors thank all participating individuals and families in these longitudinal studies.

